# High Mobility Group Box 1 enhances ADP-mediated platelet activation by increasing platelet surface P2Y_12_ localization

**DOI:** 10.1101/2021.03.24.436776

**Authors:** Deirdre Nolfi-Donegan, Gowtham K Annarapu, Lisa M Maurer, Cheryl A Hillery, Sruti Shiva

## Abstract

Thrombosis and inflammation are intimately linked and synergistically contribute to the pathogenesis of a number of vascular diseases. On a cellular level, while the platelet is central to thrombus formation as well as an active mediator of inflammation, the molecular mechanisms of cross-talk between thrombosis and inflammation remain elusive. High-Mobility Group Box 1 protein (HMGB1) is an inflammatory regulator that also stimulates platelet activation through its interaction with toll-like receptor 4 (TLR4). However, it remains unclear whether cross-talk between HMGB1 and traditional thrombotic agonists exists to modulate platelet activation. Using isolated human platelets, we tested whether HMGB1 treatment affects platelet activation mediated by traditional agonists. We found that HMGB1 enhances ADP-mediated platelet activation, but not platelet activation stimulated by thrombin or collagen. Further, inhibition of the canonical ADP purinergic P2Y_12_ receptor attenuates HMGB1-dependent platelet activation. Mechanistically, we discovered that HMGB1 activates platelet surface TLR4 to release ADP from the platelet and concomitantly increase the localization of P2Y_12_ on the platelet membrane. These data demonstrate that ADP-dependent P2Y_12_ activation contributes to HMGB1 mediated platelet activation, while HMGB1 primes platelets for an enhanced activation response to ADP. These novel findings further our understanding of thrombo-inflammatory signaling and provide new insight for therapeutic P2Y_12_ inhibition.

**Key Points:**

- HMGB1 enhances ADP-mediated platelet activation but not platelet activation stimulated by collagen or thrombin.
- HMGB1 stimulates platelet ADP release and increases platelet surface localization of P2y12 receptors via TLR4-dependent mechanism(s).

**Visual Abstract:** **Caption:** HMGB1 activates TLR4 to activate platelets, release platelet ADP, and upregulate P2Y_12_ at the platelet surface.

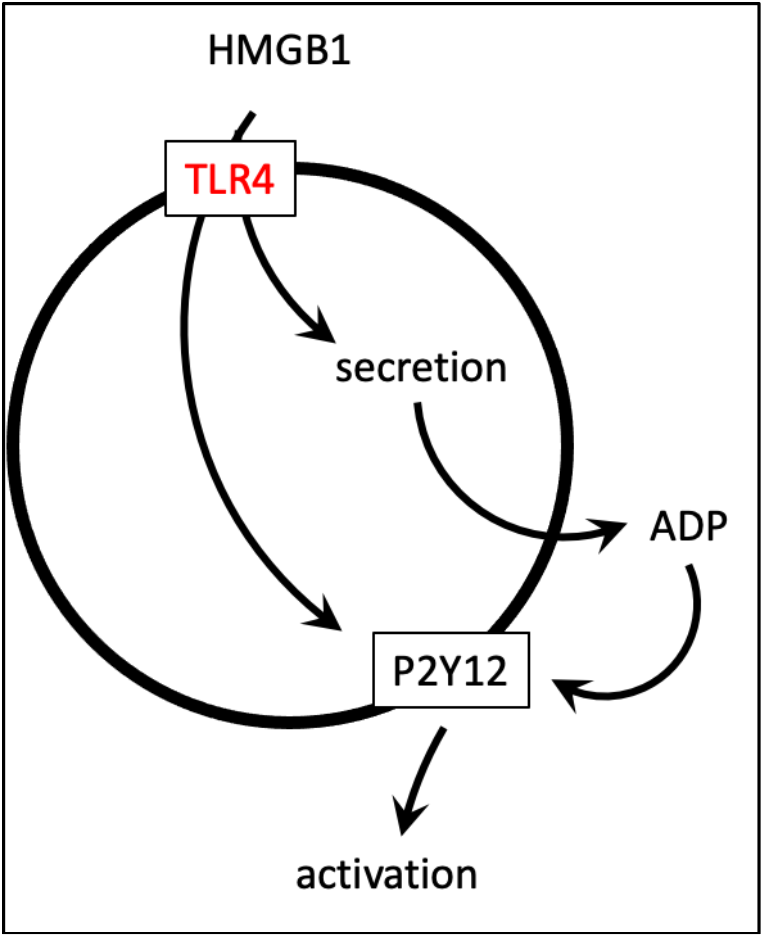

## Introduction

Inflammation and thrombosis are intimately linked, and the platelet represents a cellular mediator of thrombus formation as well as a sentinel for inflammatory signaling^1,2^. To this end, enhanced platelet activation characterizes many acute and chronic inflammatory states including diabetes^3^, sickle cell disease^4,5^, sepsis^6^, and trauma^7^, and likely contributes to the thrombotic complications observed in these conditions. Despite this recognition, the mechanisms of cross-talk between inflammatory and thrombotic stimuli that potentiate platelet activation remain elusive.

High-mobility group box 1 (HMGB1) is a highly-conserved 215 amino acid DNA-binding protein that also functions as a damage-associated molecular pattern protein to mediate sterile inflammation^8^. HMGB1 is increased in many inflammatory diseases including cancer, sepsis, sickle cell disease, autoimmune disease, and trauma^3,9–15^, which are associated with high thrombotic risk and high platelet reactivity^16–18^. Platelets, similar to other cell types, store HMGB1 and release it or present it on their surface^19^ upon activation^18,20,21,17^. Surface expression of HMGB1 enables platelet inflammatory signaling through the formation of aggregates with monocytes or neutrophils to form neutrophil extracellular traps^22^, while HMGB1 activation of toll-like receptor 4 (TLR4) on the platelet surface activates the NOD-, LRR- and pyrin domain-containing protein 3 (NLRP3) inflammasome^23^ and stimulates the production of platelet-derived microparticles^18^. In addition to its inflammatory signaling, accumulating evidence suggests HMGB1 also mediates thrombosis. For example, platelet surface HMGB1 accelerates the rate of plasminogen activation and promotes the generation of surface-bound plasmin^19,24,25^, and exogenous HMGB1 increases the speed and area of human platelet spread on collagen^17^. Consistent with this, mice with platelet-specific knockout of HMGB1 display decreased platelet activation, aggregation, and thrombus formation, while administration of recombinant HMGB1 restores their ability to form a thrombus^17,26^. On a mechanistic level, HMGB1 is redox-regulated, and depending on its oxidation state, has been shown to stimulate platelet activation predominantly through TLR4 or the receptor for advanced glycation endproducts (RAGE)^17,22,27–30^. These studies demonstrate that HMGB1 can directly stimulate platelet activation; however it remains unclear whether this inflammatory protein also acts synergistically with other traditional platelet agonists to potentiate platelet activation during inflammation.

Adenosine diphosphate (ADP), stored in platelet dense granules, is a weak platelet agonist when released. Despite this, through activation of purinergic receptors, specifically P2Y_12_, P2X_1_, and P2Y_1_, ADP is essential for mediating calcium influx and activation of the platelet along with shape change and potentiation of aggregation^31^. Inhibitors of purinergic receptors, particularly P2Y_12_, are potent inhibitors of platelet thrombotic function and are thus utilized clinically as prophylaxis in cardiovascular disease and embolic stroke^32^. Notably, several studies have implicated P2Y_12_ receptors in modulating inflammatory responses. For example, mice or patients receiving P2Y_12_ antagonists show decreased levels of inflammatory markers including circulating CD40 ligand^33–35^, interleukin-6^36^, C-reactive protein^37,38^, and platelet-leukocyte aggregates^39^. Despite this recognition, it remains unknown whether inflammatory signaling molecules, such as HMGB1 directly regulate P2Y_12_ expression or function.

Herein, we tested whether HMGB1 synergizes with traditional platelet agonists to mediate platelet activation. We demonstrate that HMGB1 potentiates platelet activation mediated by ADP. Mechanistically, HMGB1, through the activation of TLR4, upregulates P2Y_12_ at the platelet surface via dynein-dependent transport. The implications of these studies will be discussed in the context of thrombo-inflammatory signaling and therapeutic inhibition of P2Y_12_ receptors.

## Materials and Methods

### Blood draw and platelet isolation

This study was approved by the institutional review board of the University of Pittsburgh. Written informed consent was obtained from all participants. Participants were greater than 18 years old and were excluded if they were pregnant, or taking anti-platelet or anti-coagulant medications. Blood was collected in sodium citrate by standard venipuncture using a light tourniquet or no tourniquet. To isolate platelets as previously described^5^, whole blood samples were centrifuged (500*g* for 20 minutes, 25°C) to obtain platelet-rich plasma (PRP). The PRP was centrifuged (1500*g* for 10 minutes, 25°C) in the presence of PGI_2_ (1 μg/mL) to separate a platelet pellet from platelet-poor plasma (PPP). The PPP was removed, and the platelets were washed in erythrocyte lysis buffer containing PGI_2_. The platelet suspension was centrifuged a third time (1500*g* for 5 minutes, 25°C) to isolate a platelet pellet. The supernatant was removed, and the isolated platelets were resuspended in modified Tyrodes buffer (20 mmol/L HEPES, 128 mmol/L NaCl, 12 mmol/L bicarbonate, 0.4 mmol/L NaH_2_PO_2_, 5 mmol/L glucose, CaCl_2_ 3 mmol/L, 1 mmol/L MgCl_2_, 2.8 mmol/L KCl, pH 7.4). Platelet purity was confirmed by flow cytometric measurement of CD41a expression.

### Treatments of Isolated Platelets

Isolated human platelets were incubated at room temperature for 15-20 minutes with either recombinant HMGB1 (0-40 μg/ml; R&D Systems), ADP (0-5 μM; Bio/Data Corporation), collagen (0-50 μg/ml, Bio/Data Corporation), thrombin enzyme (0.1 U/ml; Sigma-Aldrich), or matched volume of platelet buffer. In select assays involving inhibition of proteins within the TLR4 signaling pathway^40^, ADP receptor signaling pathways, or cytosolic trafficking, isolated platelets were pre-treated with an inhibitor for 15-30 minutes, followed by 20 minutes of incubation with agonist. Targets for inhibition were TLR4 (anti-TLR4 IgG, 1 μg/ml; Invivogen); interleukin-1 receptor-associated kinase 1 and 4 (IRAK 1/4; IRAK 1/4 inhibitor, 0.1 μM; Tocris); TANK-binding kinase 1 (TBK1) and IκB kinase ε (IKKε; BX795, 2 μM; Invivogen); receptor for advanced glycation end products (RAGE; FPS-ZM1, 25 μM; Calbiochem); P2Y_1_ (MRS 2500, 1 nM; Tocris); P2Y_12_ (AR-C 66096, 1 μM; Tocris); cytoplasmic dynein (dynarrestin, 50 μM; Sigma-Aldrich and ciliobrevin A, 10 μM; R&D Systems); and β−tubulin (nocodazole, 2 μM; Sigma-Aldrich).

### Measurement of ADP released by platelets

ADP released into the supernatant from human platelets was quantified with an ADP ELISA Kit (Biovision ADP Colorimetric Assay Kit, #K355) according to the manufacturer’s instructions.

### Flow Cytometry Analysis

Platelet suspensions (85k cells/μL) were fixed on ice with a solution of 1% paraformaldehyde and incubated with fluorescent antibody to platelet markers CD41a (BD Biosciences #555467), activated integrin αIIbβ3 (BD Biosciences #340507), CD62P (BD Biosciences #550888), and P2Y_12_ (Biolegend #392108) for measurement by flow cytometry as previously described^5,41^. In addition to fluorescent markers, platelets were gated using forward and side scatter. Flow cytometry data were acquired using a BD-Fortessa fluorescence-activated cell sorter (FACS) Diva Software (Becton Dickinson) and analyzed using Flowjo version 10.0 software.

#### Measurement of platelet activation

Platelet suspensions were labeled with phycoerythrin (PE)-labeled mouse anti-human CD41a ab (anti-GPIIb; BD Biosciences) to gate platelets, and either FITC-labeled PAC1 ab (BD Biosciences) to detect the activation-dependent conformational change in platelet receptor integrin αIIbβ3 (GP IIb-IIIa), or with APC-labeled CD62P antibody to bind activation-dependent exposure of P-selectin.

#### Quantification of P2Y_12_ surface levels

In select assays, phycoerythrin (PE)-labeled mouse anti-human CD41a ab was used to gate platelets and FITC-labeled monoclonal P2RY_12_ antibody (anti-P2Y_12_, Biolegend) was used to detect surface expression of platelet P2Y_12_.

### Platelet P2Y_12_ Quantification by Western Blot

Western blotting was performed by standard procedure^42^ using anti-P2Y_12_ ab (Invitrogen #702516) following cell lysis of platelets. Semiquantitative analysis of images captured by chemiluminescence on a Kodak X-OMAT 2000 Processor was performed using Image Lite software.

### Statistical analysis

The unpaired parametric t-test and ANOVA were used to compare individual group samples and to compare the differences between treatments and controls. Statistical analyses were performed using Prism 9 software. P-values <0.05 were considered significant. Data are presented as mean ± standard error of the mean (SEM) unless otherwise specified.

## Results

### HMGB1 enhances ADP-dependent platelet activation

In the first series of experiments we measured the effect of recombinant HMGB1 (0-40 μg/ml) on the activation of isolated platelets by measuring surface P-selectin expression (**Figure 1A**) and activation-dependent conformation of integrin αIIbβ3 (**Figure 1B**). Consistent with prior studies, both markers showed a concentration-dependent increase, confirming that HMGB1 directly activates platelets. To determine whether HMGB1 potentiates activation stimulated by other traditional platelet agonists, we pre-treated isolated platelets with HMGB1 (10 μg/ml) before incubating with the known endogenous platelet agonists thrombin (0, 0.025, and 0.1 U/ml; **Figure 1C**), collagen (0, 5, and 50 μg/ml; **Figure 1D**), and ADP (0, 0.5, and 5 μM; **Figure 1E**). HMGB1 pre-treatment had no significant effect on collagen- or thrombin-dependent platelet activation compared to collagen or thrombin alone. However, pre-treatment with HMGB1 significantly potentiated ADP-mediated platelet activation, with HMGB1 increasing platelet activation mediated by 0.5 μM ADP by 7-fold (P=0.0023; **Figure 1E**). These data demonstrate that HMGB1 not only independently stimulates platelet activation, but also potentiates ADP-dependent platelet activation.

**Figure 1:**
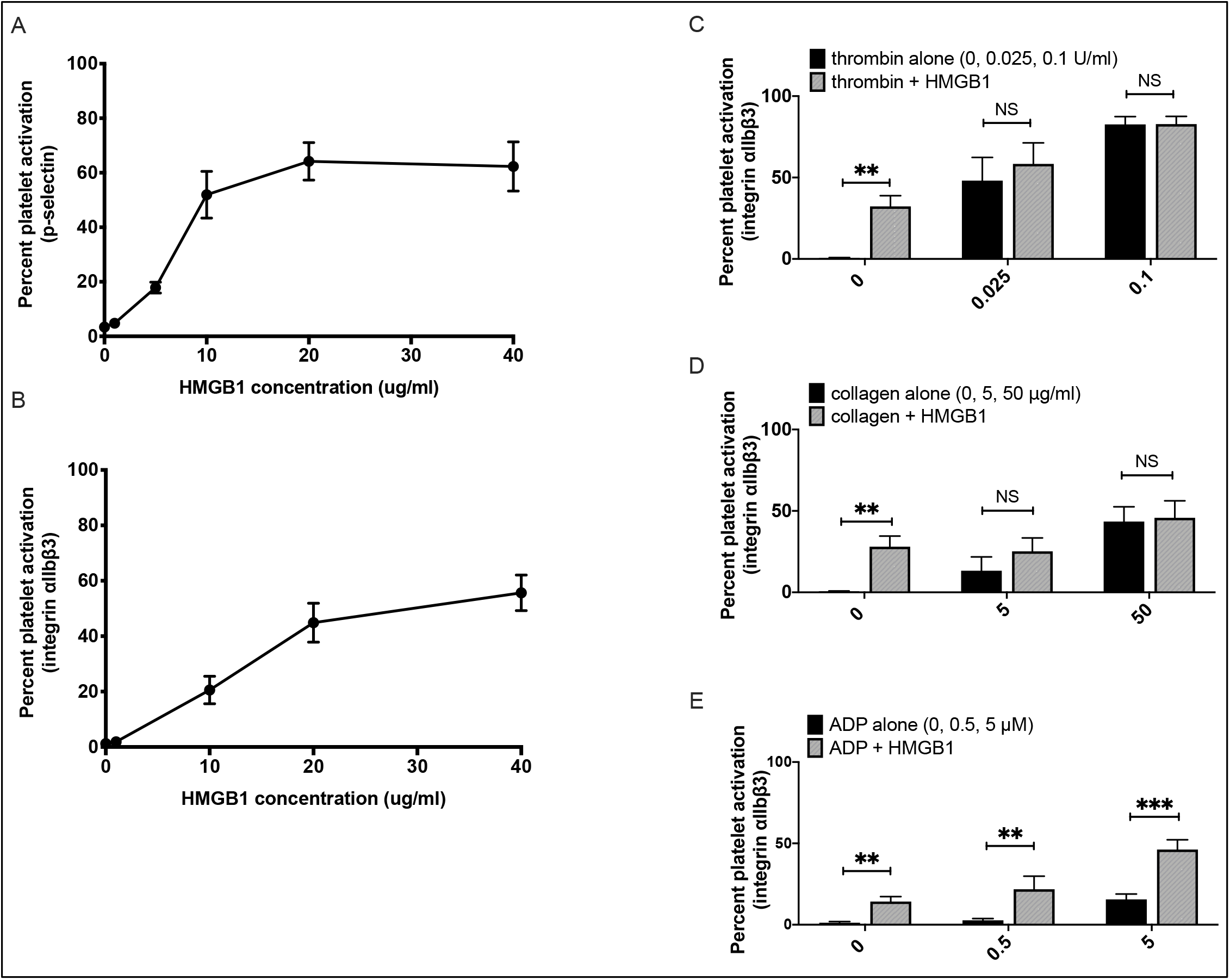
HMGB1 sensitizes platelets to ADP-dependent platelet activation. **(A-B)** Washed human platelet activation was measured by surface P-selectin expression **(A)** or integrin αIIbβ3 activation-dependent conformational change **(B)** in response to recombinant HMGB1 (0-40μg/ml; 20 min) (P<0.0001 by one-way ANOVA). Data is n=8, mean ± SEM. **(C-E)** Activation of isolated platelets treated with **(C)** collagen, **(D)** thrombin or **(E)** ADP alone (black bars) compared to the same agonist with HMGB1 pre-treatment (10 μg/ml; gray bars). n=8, data are mean ± SEM. NS = not significant,* P ≤ 0.05; ** P ≤ 0.01, *** P ≤ 0.001.

### Inhibition of TLR4 and P2Y_12_ receptor attenuates HMGB1-dependent platelet activation

To determine the mechanism by which HMGB1 potentiates ADP-dependent activation, we next tested which platelet surface receptors were required for HMGB1- and ADP-dependent platelet activation. Using pharmacologic inhibitors, we first targeted the RAGE and TLR4 receptors, which are the two predominant platelet receptors previously described to be activated by HMGB1^17,25,43,44^ (**Figure 2A**). Measurement of surface P-selectin showed that neither inhibition of RAGE by FPS-ZM1 or TLR4 using anti-TLR4 Ab had any effect on ADP-mediated platelet activation (**Figure 2B**). Inhibition of RAGE similarly had no effect on platelet activation mediated by HMGB1 alone or HMGB1+ADP together (**Figure 2B**). However, platelet activation induced by HMGB1 alone or HMGB1+ADP was significantly decreased by TLR4 inhibition (P=0.016 and P=0.0005 respectively; **Figure 2B**).

**Figure 2:**
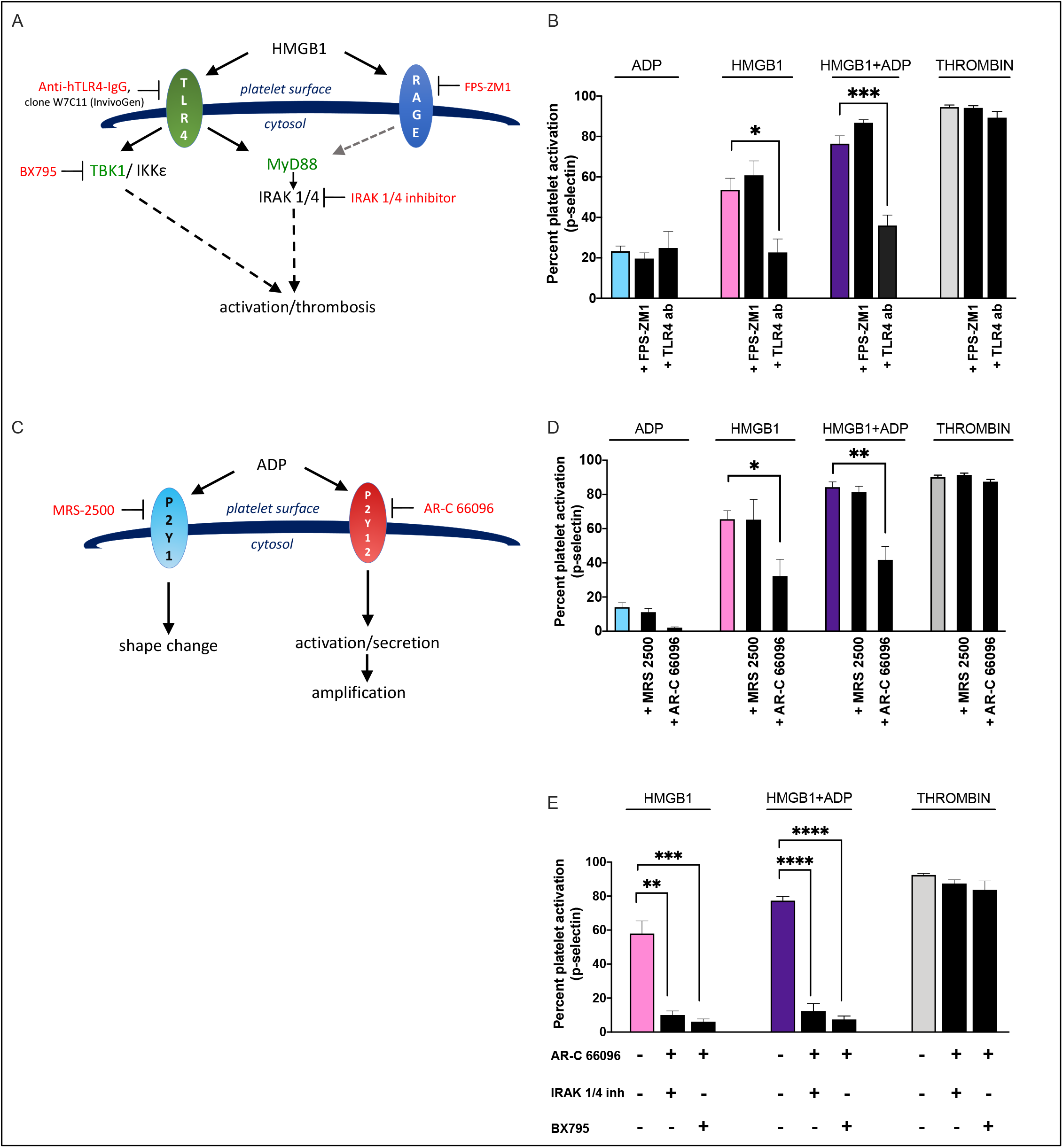
Inhibition of TLR4 and P2Y_12_ attenuates HMGB1-mediated platelet activation. **(A)** HMGB1 engages with TLR4 and RAGE whose intracellular signaling pathways converge downstream at MyD88 and IRAK 1 and 4, while MYD88-independent signaling includes TBK1. Blocking arrows depict where small-molecule inhibitors disrupt the signaling pathways. **(B)** Platelet activation stimulated with ADP (5 μM), HMGB1 (10 μM), HMGB1+ADP, or thrombin (0.1 U/ml) alone (colored bars and labeled above each bar) and after pre-treatment with FPS-ZMI (25 μM) or anti-TLR4 neutralizing antibody (1 μg/ml). n=5, data are mean ± SEM. **(C)** ADP engages P2Y_1_ and P2Y_12_ receptors. Blocking arrows depict small-molecule inhibitors MRS-2500 and AR-C 68096 antagonizing specific receptors. **(D)** Platelet activation stimulated by ADP, HMGB1, HMGB1 + ADP, or thrombin alone (colored bars and labeled above bars) or after pre-treatment with MRS 2500 (1 nM) or AR-C 66096 (1 μM). n=8, data are mean ± SEM. **(E)** Platelet activation stimulated by HMGB1, HMGB1 + ADP, or thrombin in the presence of inhibition of the P2Y_12_ receptor with AR-C 66096 (1 μM) and simultaneous inhibition of downstream TLR4 signaling with IRAK 1/4 inhibitor (0.1 μM) or BX795 (TBK1 inhibitor, 2 μM). n=4, data are mean ± SEM. NS = not significant, * P ≤ 0.05; ** P ≤ 0.01, *** P ≤ 0.001, \*\*\*\**P* ≤ 0.0001.

Since it is well-established that ADP signals through purinergic receptors^31^, we next tested the effect of MRS 2500 and AR-C66096, antagonists of the P2Y_1_ and P2Y_12_ receptors respectively, on ADP- and HMGB1-dependent platelet activation (**Figure 2C**). Platelets were pre-treated with these inhibitors and then stimulated with ADP, HMGB1, or with HMGB1+ADP (**Figure 2D**). As expected, blocking P2Y_1_, which signals platelet shape change but not sustained activation of integrin αIIbβ3 or P-selectin expression^45^, had no significant effect on platelet activation mediated by ADP (P=0.4127), HMGB1 (P=0.9798) or HMGB1+ADP (P=0.5457) (**Figure 2D**). In contrast, blocking P2Y_12_ reduced ADP-mediated platelet activation (P=0.0010) and reduced HMGB1+ADP-mediated activation by nearly 2-fold (P=0.0001) (**Figure 2D**). Surprisingly, blocking P2Y_12_ also attenuated platelet activation mediated by HMGB1 in the absence of ADP (P=0.0199). Moreover, simultaneously inhibiting both P2Y_12_ and TLR4 signaling with AR-C 66096 plus inhibitors of IRAK1/4 or TBK (the major downstream kinases activated by TLR4) caused a further reduction of HMGB1- and HMGB1+ADP-mediated platelet activation as measured by surface P-selectin expression (**Figure 2E**). Importantly, none of the inhibitors utilized had any effect on thrombin-mediated platelet activation, confirming the lack of non-specific effects of the antagonists. Similar results were observed using integrin αIIbβ3 as a marker of platelet activation (**Supplementary Figure)**. These data demonstrate that in addition to TLR4, P2Y_12_ receptors regulate HMGB1-mediated platelet activation.

### HMGB1 stimulates platelet ADP release

Platelets contain significant concentrations of ADP that may be released upon activation. Thus, we next tested whether HMGB1 stimulates ADP release from platelets, which then can bind P2Y_12_ to potentiate platelet activation. We first measured ADP concentration in the supernatant surrounding HMGB1-treated platelets, and found ADP concentration to be elevated compared with untreated platelets (**Figure 3A**). Pre-treatment of platelets with the TLR4 blocking antibody attenuated HMGB1-induced ADP release (P=0.003) (**Figure 3A).**These data indicate that HMGB1, through TLR4 activation, promotes the release of endogenous ADP from platelets.

**Figure 3:**
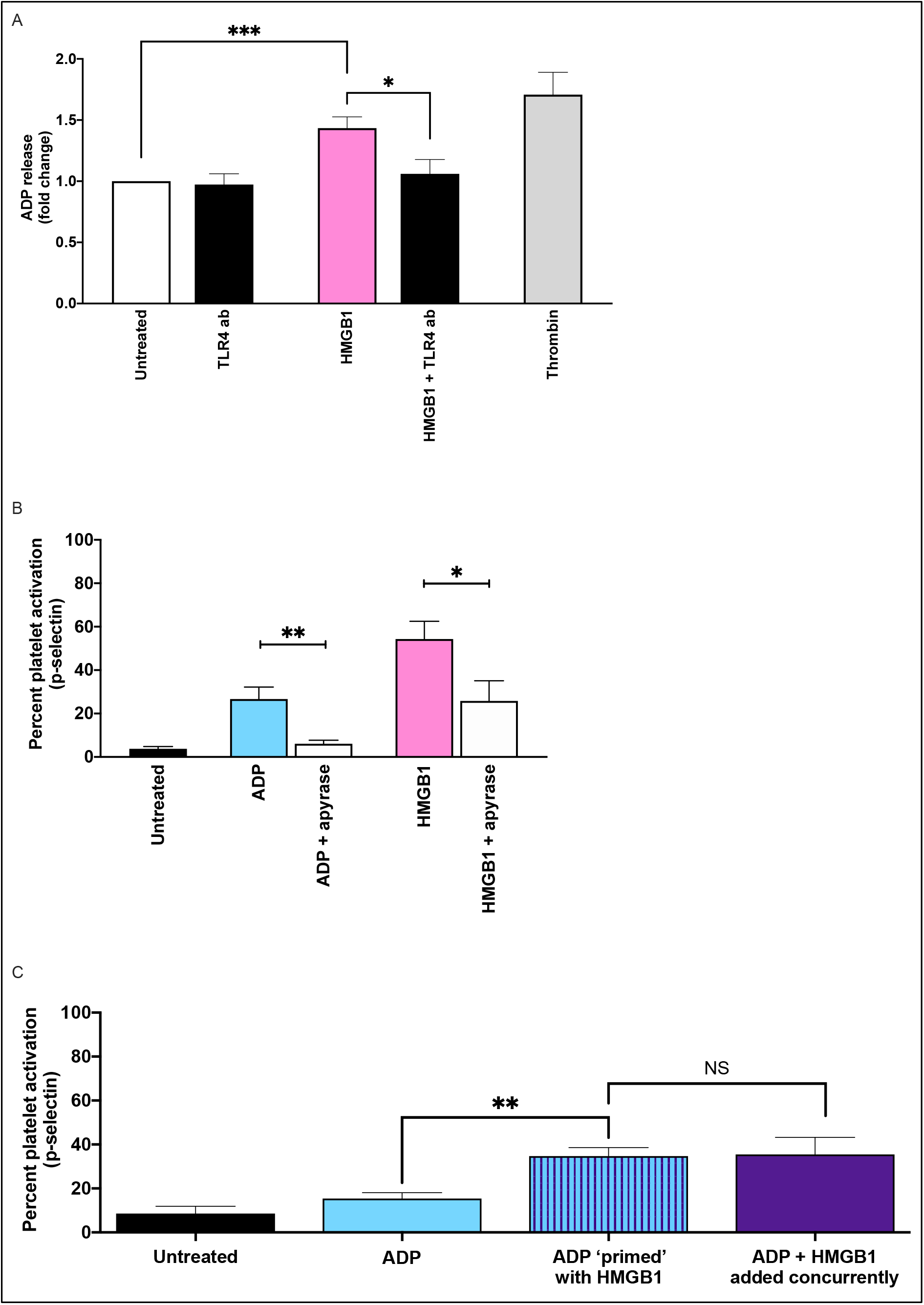
Endogenous ADP release contributes to HMGB1-dependent platelet activation. **(A)** ADP released by platelets in response to treatment with HMGB1 (10 μg/ml) and in the presence and absence of pre-treatment with TLR4 neutralizing antibody (1 μg/ml) or with thrombin (0.1 U/ml). n=4, data are mean ± SEM. **(B)** Platelet activation stimulated by ADP (5 μM, blue bar) or HMGB1 (10 μg/ml, pink bar), alone or in the presence of apyrase (0.02 U/ul; white bars). n=5, data are mean ± SEM. **(C)** Platelet activation in response to no treatment (black bar), ADP alone (5 μM, blue bar), compared to priming with HMGB1 (10 μg/ml) for 20 minutes prior to washing and adding ADP (striped bar), or treatment with HMGB1+ADP concurrently (purple bar). n=6, data are mean ± SEM. * P ≤ 0.05; ** P ≤ 0.01, *** P ≤ 0.001.

To test whether the released ADP potentiated HMGB1-mediated platelet activation, we treated platelets with HMGB1 in the presence and absence of apyrase, to degrade secreted ADP. We found that HMGB1-dependent platelet activation was attenuated by more than 2-fold in the presence of apyrase (P=0.041), suggesting that platelet-derived ADP importantly contributes to HMGB1-mediated activation (**Figure 3B**). Next, we incubated platelets with HMGB1 and then washed the platelets to remove HMGB1 and endogenous platelet-derived ADP from the suspension. Treatment of these washed platelets with exogenous ADP (5 μM) showed the same level of activation as exogenous HMGB1+ADP when added simultaneously (**Figure 3C**). Taken together, these data demonstrate that endogenous ADP release contributes to HMGB1-dependent platelet activation, and that HMGB1 also “primes” platelets for an enhanced response to ADP-stimulated activation.

### HMGB1 increases P2Y_12_ localization at the platelet surface

To determine the mechanism by which HMGB1 potentiates ADP-dependent platelet activation, we examined surface P2Y_12_ receptor levels by flow cytometry. We measured platelet surface P2Y_12_ levels before and after (20 min) treatment with ADP alone, HMGB1 alone, or HMGB1+ADP. Compared to untreated platelets, there was no significant change in surface P2Y_12_ with ADP treatment alone (P=0.44). However, stimulation with HMGB1 alone or with HMGB1+ADP resulted in a significant increase in platelet surface P2Y_12_ levels compared with untreated platelets (P<0.0001) (**Figure 4A**). Consistent with these data, HMGB1 alone induced an increase in platelet P2Y_12_ surface levels in a concentration-dependent manner (P<0.0001) (**Figure 4B**). Notably, inhibitors of the TLR4-dependent signaling molecules IRAK 1/4 (P=0.0006) and TBK1 (P=0.02) both attenuated HMGB1-stimulated increased platelet surface P2Y_12_ (**Figure 4C**).

**Figure 4:**
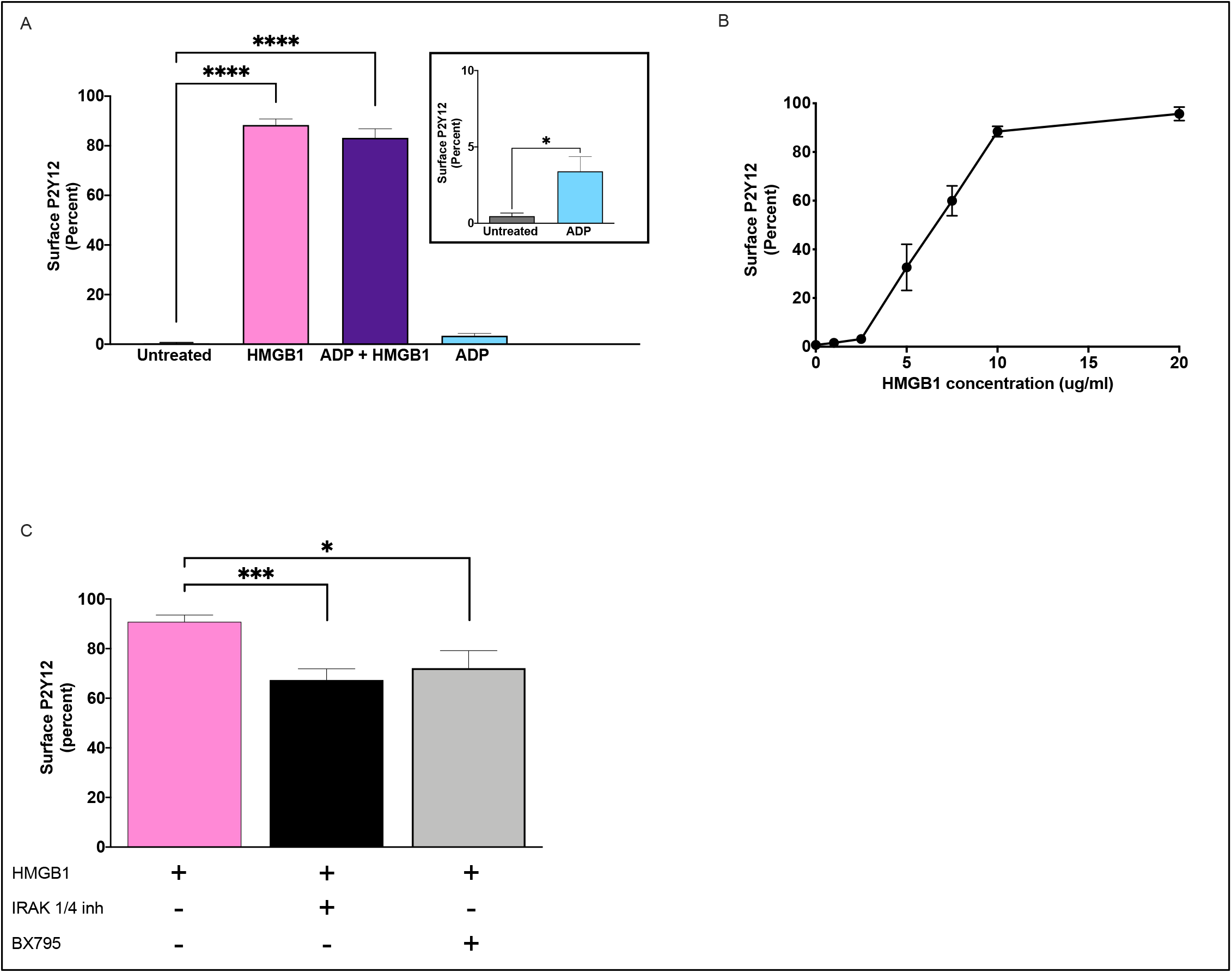
HMGB1 increases P2Y_12_ localization on the platelet surface. **(A)** Platelet surface expression of P2Y_12_ was assessed by flow cytometry after treatment with no agonist (untreated), HMGB1 (10 μg/ml), ADP alone (5 μM) (pictured inset) or HMGB1+ADP (5 μM), n=6. **(B)** P2Y_12_ surface expression in response to increasing concentrations of HMGB1 (P<0.0001 by one-way ANOVA), n=3. **(C)** Surface P2Y_12_ levels after HMGB1 (10 μg/ml) treatment in the presence and absence of IRAK 1/4 inhibitor (0.1 μM) or TBK1 inhibitor (BX795, 2 μM). n=5, data are mean ± SEM. *P ≤ 0.05; ** P ≤ 0.01, *** P ≤ 0.001.

### HMGB1-induced P2Y_12_ transportation to the platelet surface is mediated by dynein

To determine whether the HMGB1-induced P2Y_12_ surface increase was due to *de novo* protein synthesis or inhibition of proteosomal degradation, we pre-treated platelets with either cycloheximide (20 μM; 30 min) to inhibit protein translation or MG-132 (30 μM; 30 min) to inhibit protein degradation, prior to HMGB1 treatment, and then total platelet P2Y_12_ expression was measured by Western blot. HMGB1 had no effect on total platelet P2Y_12_ levels at baseline or after treatment with cycloheximide or MG-132 (**Figure 5**). These data indicate that HMGB1 does not alter whole platelet P2Y_12_ synthesis or degradation.

**Figure 5:**
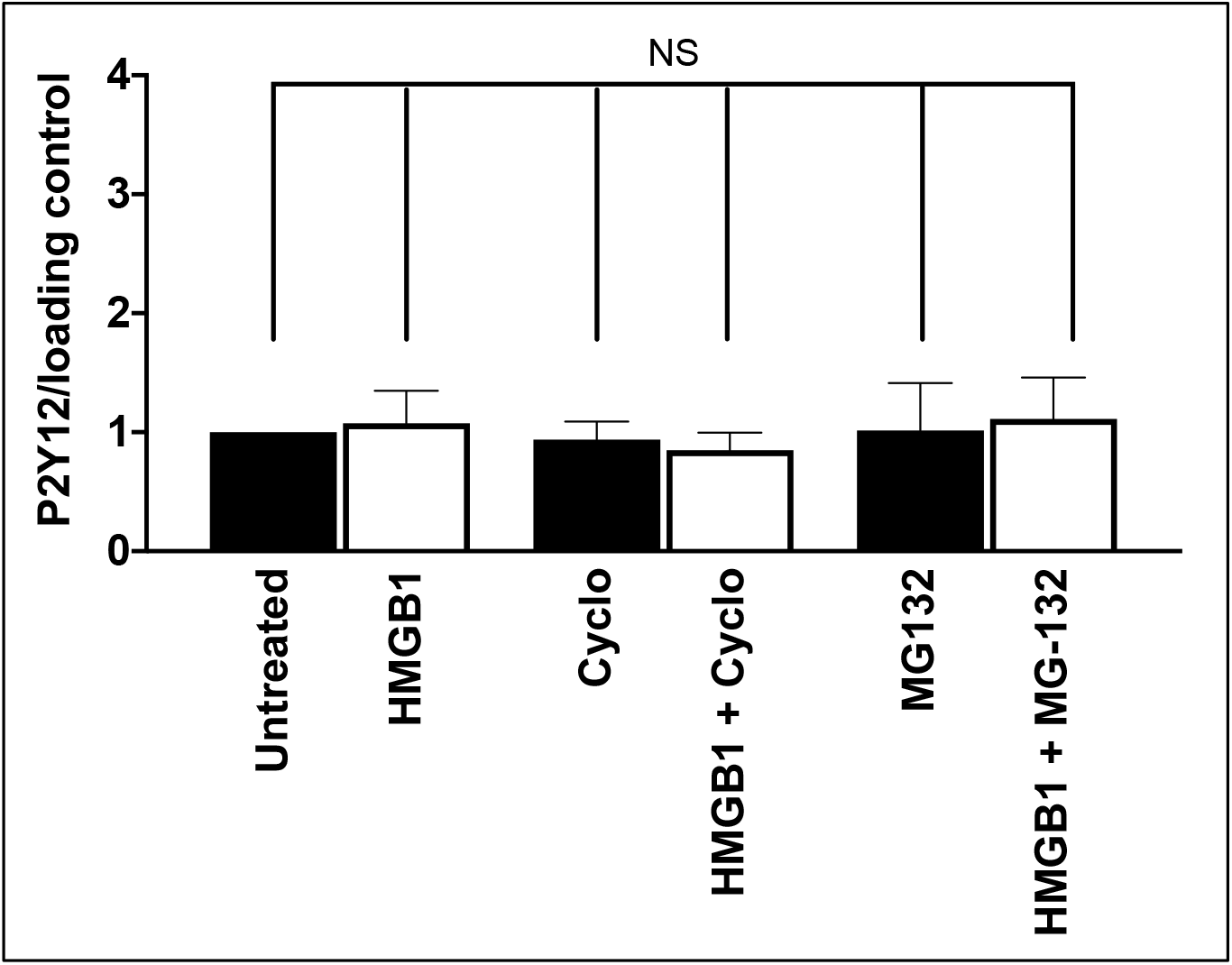
HMGB1 does not change platelet P2Y_12_ synthesis or degradation. Western blot quantification of P2Y_12_ levels in the platelet after treatment with cycloheximide (Cyclo, 20 μM) or MG-132 (30 μM) in the presence and absence of HMGB1 treatment (10 μg/ml). Total platelet P2Y_12_ levels are displayed relative to each lane’s loading control (either vinculin or GAPDH). n=4, data are mean ± SEM. * P ≤ 0.05; ** P ≤ 0.01, *** P ≤ 0.001.

Given that HMGB1 does not alter total P2Y_12_ protein synthesis, we next asked whether HMGB1 increases transport of P2Y_12_ to the platelet surface. Dynarrestin and ciliobrevin A, both small molecule inhibitors of cytoplasmic dynein, were used to disrupt dynein-mediated translocation to the surface. Pre-treatment with either dynarrestin or ciliobrevin A significantly attenuated HMGB1-dependent localization of P2Y_12_ at the platelet surface (P=0.0017 for dynarrestin, P=0.0011 for ciliobrevin) (**Figure 6A**). Similarly, treatment with nocodazole, an inhibitor of β–tubulin-associated microtubule polymerization, which is required for dynein-mediated transport, also decreased HMGB1-dependent surface P2Y_12_ levels (P=0.001) (**Figure 6A**). Measurement of platelet activation in the presence of the inhibition of cytoplasmic dynein or microtubule polymerization had no effect on ADP-mediated platelet activation. However, inhibition of P2Y_12_ transport significantly attenuated platelet activation stimulated by HMGB1 alone and abolished the HMGB1-mediated sensitization of platelets to ADP (**Figure 6B-C**). These data demonstrate that HMGB1 promotes dynein-dependent mobilization of P2Y_12_ from internal stores to the platelet surface, and that dynein-dependent transport not only contributes to HMGB1-induced platelet activation, but also underlies the HMGB1-mediated sensitization of platelets to ADP.

**Figure 6:**
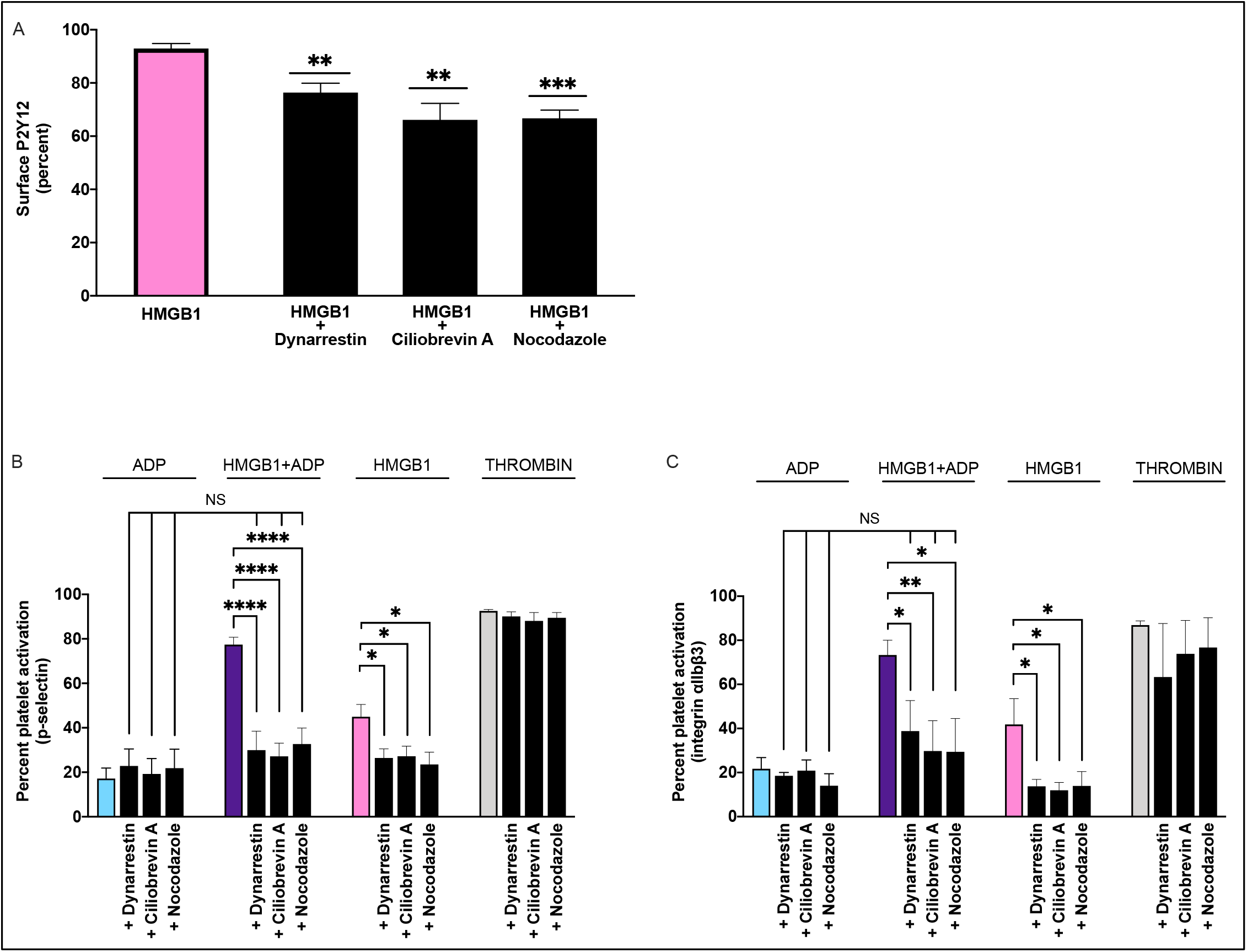
Cytoplasmic trafficking inhibitors attenuate surface P2Y_12_ expression and priming effect of HMGB1. **(A)** Platelet P2Y_12_ surface levels (n=5) and platelet activation (n=4) measured by **(B)** surface P-selectin expression or **(C)** integrin αIIbβ3 in response to pre-treating platelets with dynarrestin (50 μM), ciliobrevin A (10 μM), or nocodazole (2 μM) prior to HMGB1 treatment (10 μg/ml). Data are mean ± SEM.* P ≤ 0.05; ** P ≤ 0.01, *** P ≤ 0.001.

## Discussion

In this study, we show for the first time that HMGB1 and ADP synergize to enhance platelet activation. HMGB1-dependent TLR4 activation releases endogenous ADP from the platelet, which potentiates the platelet activation mediated by HMGB1. Concomitantly, HMGB1 increases transport of P2Y_12_ receptors to the platelet surface, which “primes” the platelet for an enhanced activation response to ADP. These findings illustrate a novel mechanism by which P2Y_12_ receptors are involved in inflammatory signaling, as well as a new line of synergy between the inflammatory and thrombotic pathways.

Our study confirms prior work demonstrating that HMGB1 induces platelet activation in a TLR4-dependent manner^17,23,29,30^. We further demonstrate that this activation is partially dependent on TLR4-mediated ADP release and P2Y_12_ activation. These data put HMGB1 in the ranks of other platelet agonists such as thrombin and collagen, which stimulate release of endogenous platelet-derived ADP to fuel feed-forward platelet activation and aggregation through P2Y_12_^46,47^ Although ADP is considered a relatively weak physiologic agonist, it is essential for recruiting and cross-linking additional platelets^45,48,49^ and stabilizing the outer shell of the platelet plug^50^. While prior studies have shown that lipopolysaccharide (LPS), a strong TLR4 agonist, induces platelet dense and α-granule release^51^, P2Y_12_ inhibition has shown mixed results on LPS-induced platelet activation and coagulation^52–55^. Our data showing an approximately 50% decrease in HMGB1-induced platelet activation in the presence of P2Y_12_ inhibition may serve as a new foundation for studies investigating differences in LPS and HMGB1-mediated platelet signaling.

Our data demonstrate that HMGB1 primes platelets to ADP-induced activation; this is consistent with a prior study in which HMGB1 caused increased sensitivity to ADP-stimulated platelet aggregation^22^. We extend these studies to demonstrate that HMGB1-stimulated P2Y_12_ receptor transport mechanistically underlies this priming effect. Human platelets express P2Y_12_ both on their surface and in an inducible pool within the platelet that can be rapidly mobilized^56^. It is estimated that about half the quantity of platelet P2Y_12_ is found on the plasma surface membrane and in open canalicular system at all times^57^, but the mechanisms that regulate platelet P2Y_12_ surface expression are not fully elucidated^58^. Here we show that HMGB1 stimulates transport of P2Y_12_ to the platelet surface, and this transport is partially dependent on TLR4 signaling and dynein activity. While it has been demonstrated that LPS increases expression of P2Y_12_ on macrophages^59^, this type of TLR4-dependent regulation of the purinergic receptor has not been described in the platelet or with any other TLR4 ligand. Further study is required to delineate the molecular link between TLR4 activation and the regulation of dyneins. Of note, phosphorylation and translocation of these motor proteins within the platelet has been shown to occur after activation^60^, suggesting that perhaps TLR4-dependent phosphorylation may play a role in P2Y_12_ translocation.

Beyond thrombotic signaling, our data provide a potential mechanistic link for the growing number of studies that implicate P2Y_12_ in inflammatory signaling^61,62^. The platelet is an established sentinel of inflammatory signaling that not only synthesizes, stores, and releases chemokines and cytokines, but also interacts with and activates leukocytes to propagate inflammatory signaling^63^. Notably, genetic or pharmacologic inhibition of P2Y_12_ has been shown in both murine models and human studies to decrease levels of circulating inflammatory mediators such as TNF-α, IL-10, IL-6 and MIP-1β,^62^ and attenuate the release of soluble CD40 ligand^33^. Further, P2Y12 inhibition has attenuated platelet-neutrophil aggregates and neutrophil extracellular trap formation in multiple models of LPS-induced inflammation^64–66^. Consistent with this, P2Y_12_ inhibition has shown beneficial effects in clinical studies of patients with inflammatory syndromes including pneumonia and sepsis^61,67^. Mechanisms underlying the P2Y_12_-mediated attenuation of inflammation remain elusive, and have been predominantly investigated in LPS-induced inflammatory models^64,66^. Our data showing HMGB1-induced ADP release and increased P2Y_12_ surface expression under the control of TLR4 provide a novel mechanism of P2Y_12_ regulation of platelet inflammatory signaling. Additionally, given that LPS can induce the release of HMGB1 to potentiate inflammatory signaling, the data presented here provide a rationale to delineate the cross-talk between different TLR4 agonists and ADP-dependent signaling. Moreover, these data suggest that P2Y_12_ activity and expression potentially play a significant role in the propagation of sterile inflammation.

Pharmacologic inhibitors of P2Y_12_ are widely used therapeutically for a number of pathologies ranging from cardiovascular diseases to sepsis, and rates of resistance to therapeutic P2Y_12_ inhibition have been reported as high as 30% in patients treated with clopidogrel^68^. It is interesting to consider whether HMGB1-induced P2Y_12_ surface localization potentially underlies resistance, particularly in patients with inflammatory disease. Consistent with this idea, Haberstock-Debic et al^69^ previously showed that platelet stimulation with strong agonists such as thrombin significantly increase surface P2Y_12_ expression, and this increased expression was associated with resistance to clopidogrel and could be reversed by elinogrel, a P2Y_12_ inhibitor with longer half-life which was able to antagonize the higher levels of receptors expressed. In pathologies with inflammatory components, acute HMGB1 release from the platelet or other cells could potentially confer resistance to purinergic antagonists. However, further studies are required to dissect this signaling pathway both mechanistically and in patients treated with P2Y_12_ inhibitors.

In summary, this study shows for the first time that ADP release and P2Y_12_ activation contributes to HMGB1-induced platelet activation, and that HMGB1 primes platelets for ADP signaling by increasing P2Y_12_ at the platelet surface without invoking new P2Y_12_ synthesis. These data present a novel mechanism of cross-talk between inflammatory and thrombotic signaling and have implications for both the therapeutic targeting of HMGB1 and P2Y_12_ receptors.

## ACKNOWLEDGEMENTS

This work was supported by NIH grants HL130268-03 (to SS), HL128371 (to CAH), and HD071834 (to DND). Additional support was provided by The Hemophilia Center of Western Pennsylvania (to SS), the UPMC Children’s Trust Young Investigator Foundation Fund 020284 (to DND), and the Children’s Hospital of Pittsburgh Scholars Award (to DND).

## AUTHORSHIP CONTRIBUTIONS

DND and SS designed the study. DND performed the research. DND, GKA, LMM, SS, and CAH analyzed the data. DND, SS, and CAH wrote the manuscript.

## Supplementary Figure

**Supplementary Figure:**
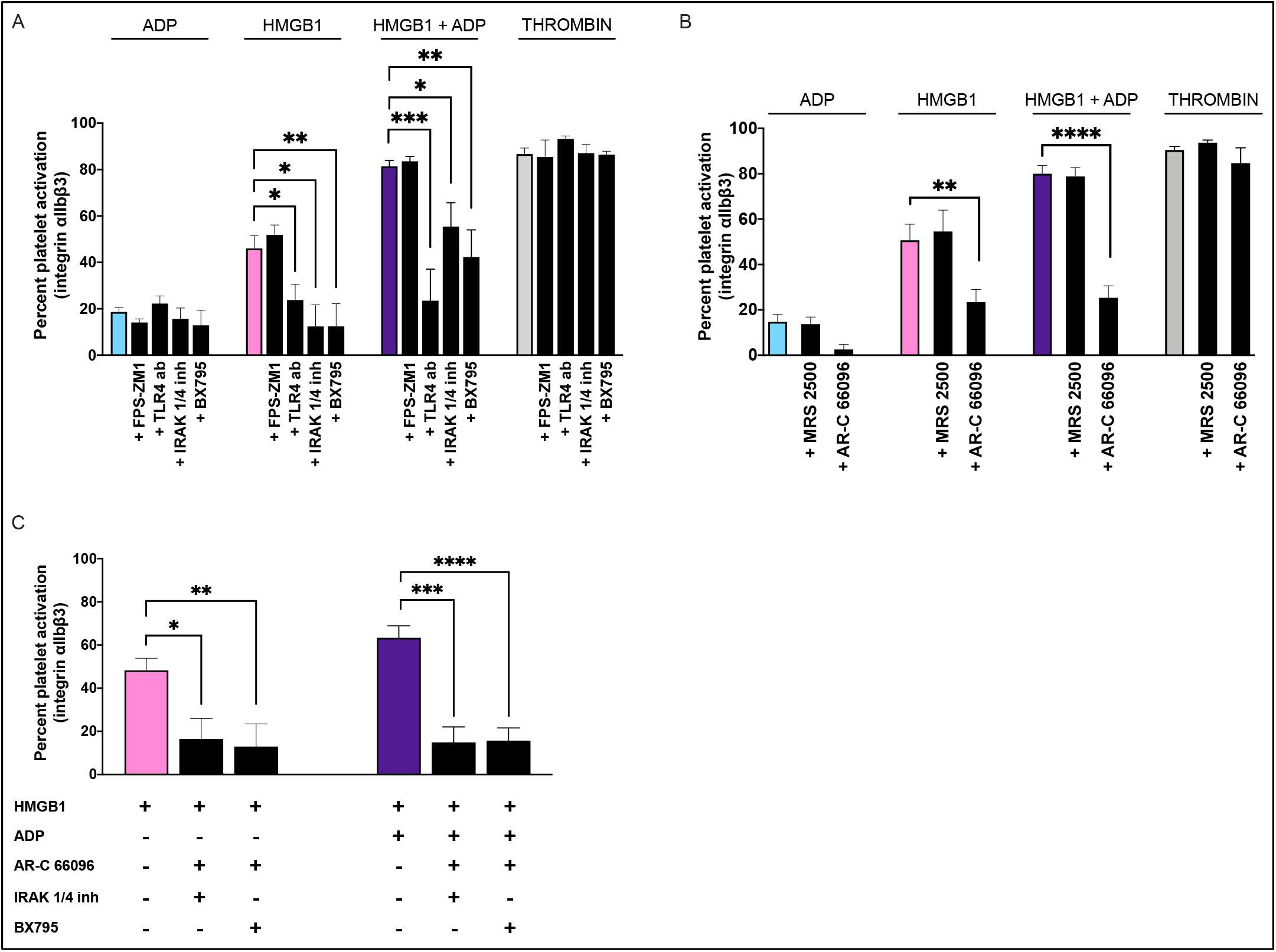
TLR4 and purinergic receptor inhibitors attenuate HMGB1-dependent activated integrin αIIbβ3 in human platelets. **(A)** Platelet activation stimulated with HMGB1 alone (10 μg/ml), ADP alone (5 μM), HMGB1+ADP, or thrombin alone and after pre-treatment with FPS-ZMI (25 μM), anti-TLR4 neutralizing antibody (1 μg/ml), IRAK 1/4 inhibitor (0.1 μM) or BX795 (2 μM). n=3. **(B)** Platelet activation stimulated by ADP, HMGB1, HMGB1 + ADP, or thrombin alone (colored bars) or after pre-treatment with MRS 2500 (1 nM) or AR-C 66096 (1 μM). n=4, data are mean ± SEM. **(C)** Platelet activation stimulated by HMGB1 alone (pink bar) or HMGB1+ADP (purple bar) while antagonizing the P2Y_12_ receptor with AR-C 66096 (1 μM) while simultaneously inhibiting TLR4 signaling with IRAK 1/4 inhibitor (0.1 μM) or BX795 (TBK1 inhibitor, 2 μM). n=4, data are mean ± SEM.* P ≤ 0.05; ** P ≤ 0.01, *** P ≤ 0.001.

